# Genetic bases of resistance to the rice *hoja blanca* disease deciphered by a QTL approach

**DOI:** 10.1101/2022.11.07.515427

**Authors:** Alexander Silva, María Elker Montoya, Constanza Quintero, Juan Cuasquer, Joe Tohme, Eduardo Graterol, Maribel Cruz, Mathias Lorieux

## Abstract

Rice *hoja blanca* (RHB) is one of the most serious diseases in rice growing areas in tropical Americas. Its causal agent is *Rice hoja blanca virus* (RHBV), transmitted by the planthopper *Tagosodes orizicolus* Müir. Genetic resistance is the most effective and environment-friendly way of controlling the disease. So far, only one major quantitative trait locus (QTL) of *Oryza sativa* ssp. *japonica* origin, *qHBV4.1*, that alters incidence of the virus symptoms in two Colombian cultivars has been reported. This resistance has already started to be broken, stressing the urgent need for diversifying the resistance sources. In the present study we performed a search for new QTLs of *O. sativa indica* origin associated with RHB resistance. We used four F_2:3_ segregating populations derived from *indica* resistant varieties crossed with a highly susceptible *japonica* pivot parent. Beside the standard method for measuring disease incidence, we developed a new method based on computer-assisted image processing to determine the affected leaf area (ALA) as a measure of symptom severity. Based on the disease severity and incidence scores in the F_3_ families under greenhouse conditions, and SNP genotyping of the F_2_ individuals, we identified four new *indica* QTLs for RHB resistance on rice chromosomes 4, 6 and 11, namely *qHBV4.2^WAS208^*, *qHBV6.1^PTB25^*, *qHBV11.1* and *qHBV11.2*. We also confirmed the wide-range action of *qHBV4.1*. Among the five QTLs, *qHBV4.1* and *qHBV11.1* had the largest effects on incidence and severity, respectively. These results provide a more complete understanding of the genetic bases of RHBV resistance in the cultivated rice gene pool, and can be used to develop marker-aided breeding strategies to improve RHB resistance. The power of joint-and meta-analyses allowed precise mapping and candidate gene identification, providing the basis for positional cloning of the two major QTLs *qHBV4.1* and *qHBV11.1*.

## Introduction

The rice *hoja blanca* (RHB) disease is one of the most important constraints to rice productivity in the tropical zone of Americas, causing yield losses in many countries, including Colombia, Costa Rica, Ecuador, Guyana, Panama, Peru, Dominican Republic, Nicaragua and Venezuela (Morales and Jennings 2010). Its causal agent is the *Rice hoja blanca virus* (RHBV), a *Tenuivirus* transmitted by the planthopper *Tagosodes orizicolus* M. (Hemiptera:Delphacidae). Genetic resistance to both the virus and its vector insect is key for a successful, environment-and consumers health-friendly, integrated crop management. No real immunity has been found in the cultivated rice germplasm, and even in resistant materials, the plantlets (< 10 days old) can show susceptibility to the virus. Nonetheless, a handful of varieties – including the two Colombian cultivars, Fedearroz 50 and Fedearroz 2000 – with good resistance level have been bred in the past and have been important to stabilize rice production. Fedearroz 2000 is still the most resistant commercial variety, however, the incidence of the disease has increased under field and controlled conditions. Fedearroz 50 has turned virus-susceptible in the past years in the field, probably due to virus mutations that allowed it to overcome the resistance. Thus, there is an urgent need for diversifying the sources of genetic resistance in order to breed new varieties with more durable resistance. Genetic resistance to RHB disease can be decomposed into resistance to the virus itself, and resistance to its insect vector.

In a previous study, we reported a major QTL on the short arm of chromosome 4 for resistance to RHBV, shared by Fedearroz 50 and Fedearroz 2000, abbreviated as FD 50 and FD 2000 in this paper (Romero et al. 2014). This QTL, called here *qHBV4.1*, controlled RHB *incidence*, measured as the percentage of plants that show symptoms of the viral infection, no matter the level of the symptoms. Incidence is of course an important parameter of the epidemics of a disease. Yet, its *severity* is certainly as much as important: if severity is low, a high incidence might have no significant impact on plant viability, panicle development, or grain yield. Additional to RHBV incidence, we thus designed – and report in this study – new experiments to decipher the genetic control of RHB resistance seen as symptoms severity, measured by computer-aided image processing of the affected leaf area (ALA).

The *Oryza sativa* species is divided into two main clades, *indica* and *japonica*, which can be considered as sub-species. The *indica* and *japonica* are themselves divided into 15 subgroups (Li et al. 2014). Using genomic sequencing data, it is possible to infer the *indica* or *japonica* origin of an accession at a particular chromosome location by local ancestry analysis (Santos et al. 2019). We showed by local ancestry analysis showed that, although FD 50 and FD 2000 are mostly of *indica* genetic background, *qHBV4.1* in those two cultivars was found to be of *japonica* origin and was surrounded by several hundred kilobase pairs of *japonica* DNA. We found desirable to search for *indica* resistance QTLs in order to ensure better compatibility with tropical irrigated materials in breeding programs, and also to increase the diversity of the available sources of genetic resistance. Based on the encouraging screening work by the FLAR (Fondo Latinoamericano para Arroz de Riego) team (Cruz-Gallego et al. 2018), we thus designed a new QTL analysis based on four crosses involving new resistance sources.

## Materials and methods

### Plant materials

Four *O. sativa* ssp. *indica* accessions were selected amongst the most resistant materials identified in (Cruz-Gallego et al. 2018): WAS 208-B-B-5-1-1-3 (IRGC 121855) thereafter abbreviated as WAS 208), Badkalamkati (IRGC 45011) (abb. Badka), PTB 25 (IRGC 6386) and Fedearroz 2000 (IRGC 124388) (abb. FD 2000). The main selection criteria were (1) selected sources should show consistent and high resistance to RHB and (2) they should cover the genetic diversity spectrum of the *indica* cluster. Then, the four resistance sources were crossed with the highly susceptible *japonica* line Bluebonnet 50 (IRGC 121874) (abb. BBT 50), using BBT 50 as the male parent. Candidate F1 hybrids were checked for self-pollination of the mother plant with 48 SNP (single nucleotide polymorphism) markers distributed along the rice genome. F_2_ (S_1_) populations were derived from self-pollination of the verified F_1_ hybrids. Each F_2_ plant was self-pollinated to produce F_3_ (S_2_) families. All panicles involved in crosses or self-pollination were bagged to avoid out-crossing.

### Population sizes

The following population sizes were used for both genotyping in F_2_ plants and trait scoring in F_3_ families: WAS 208 × BBT 50: 104 F_2_, 1,040 F_3_ (ALA), 6,240 F_3_ (incidence); Badka × BBT 50: 105 F_2_, 1,050 F_3_ (ALA), 6,300 F_3_ (incidence); PTB 25 × BBT 50: 108 F_2_, 1,080 F_3_, 6,480 (incidence); FD 2000 × BBT 50: 105 F_2_, 1,050 F_3_ (ALA), 6,300 F_3_ (incidence). Populations sizes for ALA scoring are significantly lower than for incidence, however, as ALA is assessed using tube-based phenotyping (non-choice feeding test), precision is higher than for incidence measured in trays (see hereby).

### Evaluation of RHB resistance

To evaluate the level of resistance – or susceptibility – of the F_3_ plants, we looked at two complementary aspects of the disease: the *incidence* and the *severity* of the symptoms. *Incidence* is simply defined by the proportion of diseased plants in a population, while *severity* is the area or volume of plant tissue that is visibly diseased (Campbell and Neher 1994). The F_3_ families, parents and check lines were evaluated under greenhouse conditions, at the CIAT (Centro Internacional de Agricultura Tropical, International Centre for Tropical Agriculture) headquarters in Palmira, Colombia.

### Virulent insect colonies

*T. orizicolus* virulent insects were obtained from colonies maintained at CIAT. The RHBV-harboring colony contained insects that were fed on RHBV-infected plants and allowed to reproduce on BBT 50. To determine the percentage of virulent insects in this colony, 200 individual nymphs were tested for virulence on separate, caged, RHB-susceptible 8-day-old seedlings.

### Incidence

Here we consider incidence as the percentage of plants showing any level of RHB symptoms. We consider the absence of symptoms as a sign of absence of infection, not as extreme tolerance. F_3_ materials and controls were planted in plastic trays containing 17 furrows of 20 plants each. Each tray contained a furrow of each parent – BBT 50 also served as a susceptible control – and the controls FD 2000 (resistant) and Colombia 1 (intermediate). The trays were placed in a mesh cage 18 days after sowing, and infested by mass release of *T. orizicolus* with virulence between 50% and 65%, with an average of four nymphs per plant. The nymphs were allowed to feed for three days on the plants, after which they were eliminated with water rinsing. A randomized complete block design was used with three replicates (20 plants from each F3 family per replicate), where each block represented a cage. Incidence assessment was made 35 days after infestation (DAI) by counting the plants showing disease symptoms, per row.

### Severity

In order to test the severity of RHB, 10 plants per F_3_ family were measured individually. 18 days after sowing, plants were transferred in individual transparent tubes with 3 nymphs per plant. Nymphs were allowed to feed for three days on the plants. Symptoms severity was measured using a novel methodology based on computer-assisted image processing. Briefly, images of the three youngest leaves infected plants were taken using a reference scale (a ruler) and a contrasting background under homogeneous light exposure. Raw images of plants with symptoms (Figure 1A) were processed using the ImageJ software (https://imagej.nih.gov/ij) in two steps: in the first step, brightness was decreased to eliminate the areas affected by the disease (Figure 1B). Then, the image was binarized, *i.e.*, converted to black and white (Figure 1C). The number of black pixels was thus representing the healthy area *H* of the leaf, according to the previously calibrated reference scale. In the second step, a binarization process allowed to calculate the total area *T* (Figure 1D). The affected leaf area (ALA) by the virus was then calculated in each F_3_ plant as:

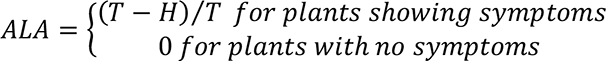

**Figure 1.**
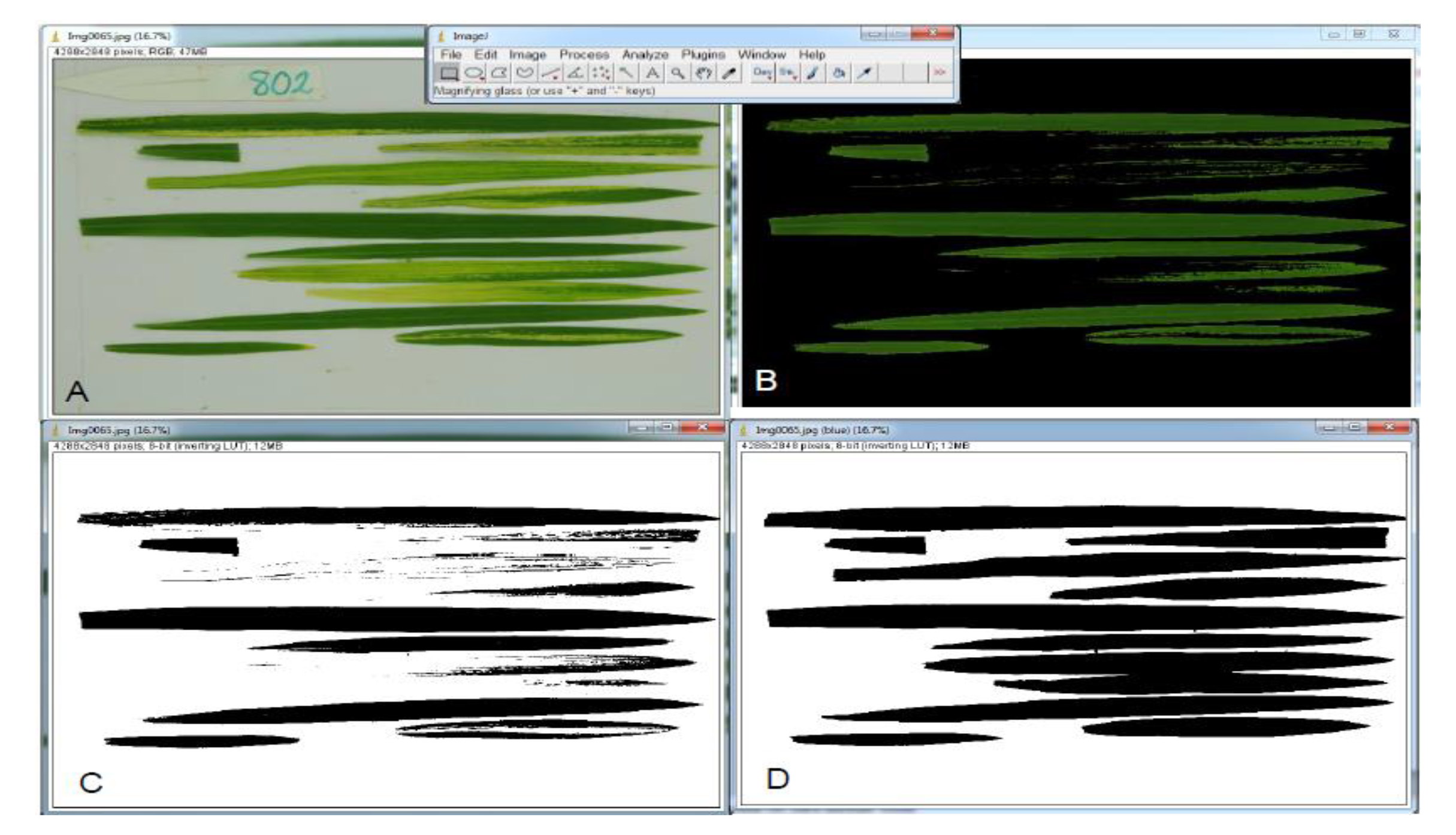
RHB disease severity (% of affected leaf area, ALA) measurement workflow. A: Image acquisition of rice leaves affected by RHB. B: Image segmentation to separate healthy from affected leaf areas. C: Healthy leaf area binarization. D: Total leaf area binarization

The ALA of each F_3_ family was then calculated as the average of the ALA values of the 10 F_3_ plants. In an attempt to remove possible correlation effects with incidence, we also calculated the averaged ALA only over the plants that showed symptoms (ALA_strict).

### Statistical treatment

Since ALA was measured as the ratio of affected/total leaf area for each family in each block, a weighted ALA was calculated with a generalized linear mixed model (MLGM) using the gamma distribution. The incidence variable was analyzed with an MLGM using the binomial distribution. The MLGM procedure is used for variables that are not necessarily normally distributed. For each variable, an analysis of variance was performed to test for differences between families and to evaluate the effect of the blocks. Subsequently, adjusted means were calculated for each family and a mean comparison test (t-test) was performed to identify those families that were more resistant or equal to the resistant parent. The descriptive analysis was performed in the R program and the other analyses were performed in the SAS^®^ statistical program.

### SNP marker development

Whole-genome sequences (WGS) of parental lines, available from previous studies (Li et al. 2014; Duitama et al. 2015; Cruz-Gallego et al. 2018) and the IRGSP 1.0 Nipponbare reference genome were compared *in silico* using the Next Generation Sequencing Eclipse Plugin, NGSEP (Duitama et al. 2014). Variant detection and annotation were performed with NGSEP and SNP positions were determined relative to the reference genome. Informative biallelic SNPs were filtered based on the following characteristics of their flanking sequence: minimum 30 bp between the target SNP and other variant and GC content between 20 and 65%. The flanking sequence of 100bp upstream and downstream the target SNP was retrieved and submitted to D3 assay design web-based tool (https://d3.fluidigm.com) to develop Fluidigm^®^ SNPtype assays. SNPs were chosen based on their design quality score and their genomic position.

### DNA extraction

Leaf tissue was collected from 15-days-old F_2_ plants. Samples were frozen in liquid nitrogen and stored at –80°C until processed. Plant DNA was isolated in 96-racked tubes using a modified version of a method previously described (Risterucci et al. 2000) as follows: 480 µl of extraction buffer was added to 150 mg of ground frozen leaf tissue. The buffer was 100 mM Tris (pH = 8.0), 2M NaCl, 20 mM EDTA (pH = 8.0), 2% MATAB, 0.5% sodium bisulfite, and 1% PEG 8000. This mixture was homogenized and incubated in a water bath at 74°C for 30 min. Subsequently, 480 µl of chloroform:isoamyl-alcohol (24:1) was added and the mixture was centrifuged at 3000 rpm in a Sorvall^®^ centrifuge with a GLA-3000 rotor. Supernatants were precipitated with 270 µl of isopropanol at –20°C for one hour and centrifuged at 3000 rpm. The pellets were washed with 200 µl of 80% ethanol and allowed to dry at 40°C by inverting the tubes for one hour. DNA was resuspended in Tris-EDTA [10 mM Tris-HCl, pH8 and 1 mM EDTA, pH8) containing 40µg/ml of RNAse A and spectrophotometric quantification was done using a multimode plate reader Synergy H1 (Biotek^®^). DNAs were normalized at 60ng/µl for subsequent processing.

### Genotype scoring

F_2_ individuals and parents were scored for polymorphic SNPs in each population using the Fluidigm^®^ nanofluidic genotyping platform, according to the manufacturer’s protocols. Briefly, pre-amplification of targets regions was done in a 5µl multiplex-PCR mixture. The amplicons were used as template for allele specific-PCR that was conducted in 48 × 48 integrated fluidic circuit (IFCs) of the Fluidigm^®^ genotyping platform. Fluorescent images of the IFCs were acquired in an EP1 reader and analyzed with the Fluidigm^®^ Genotyping Analysis Software.

### Linkage maps construction

Genetic maps were calculated with MapDisto 2.0 (Lorieux 2012; Heffelfinger et al. 2017) (http://mapdisto.free.fr). Goodness-of-fit to Mendelian segregation (1:2:1) was tested for each marker by computing the chi-squared (*χ*^2^) statistic with the ‘Segregation *χ*^2^s’ function. As the SNP markers were defined from WGS, we kept their order on the physical map or rice. For each cross separately, we checked the data for singletons (e.g., the “B” in “AAAAAAA**B**AAAAAA” is a singleton) with the ‘Replace errors by flanking genotypes’ function, with a maximum probability of a singleton to occur of 0.001. A few missing data were inferred using the ‘Replace missing data by flanking genotypes’ function, with the same threshold. Recombination fractions were calculated with the standard EM algorithm (appendix 1 in Lorieux 2021) converted to centimorgans (cM) with the Kosambi mapping function (Kosambi 1944).

### QTL mapping

Data files were prepared using the ‘Export map and data’ function of MapDisto 2.0. Analyses of distribution of the phenotypic traits as well as QTL detection were performed using the Qgene 4.0 program (Joehanes and Nelson 2008) (http://www.qgene.org). For QTL detection, the LOD score statistics were calculated with different methods and then compared: single-marker regression (SMR), simple interval mapping (SIM) and composite interval mapping (CIM). The forward cofactor selection option was used in CIM. Additivity and dominance effects were calculated according to Falconer (1960). Empirical thresholds to declare the presence of a QTL were obtained using the resampling by permutation method, performing 10,000 (SMR) or 1,000 (CIM) iterations for each trait-chromosome combination. In order to correct for possible erroneous phenotypic data corresponding to escape, mis-scoring, or incomplete penetrance, all positive QTLs were additionally confirmed by analysis of outliers in the trait distribution using the ‘Plot trait vs. genotype’ module of MapDisto. This module allows to calculate corrected single-marker regression F-test values after detecting and removing outlier data in each marker genotypic sub-class.

In the case of a secondary LOD score peak linked to a major peak of a QTL, in order to determine if the secondary peak corresponded to a true QTL or to an artifact – or “fake QTL” – a detailed analysis of the distribution of recombination fractions along the chromosome was performed, following the method of Lorieux (2018). The analysis looked for restriction of recombination fractions that could induce artificial linkage disequilibrium between the major and the secondary LOD score peaks. If artificial linkage disequilibrium was detected, then the secondary peak was declared an artifact. Interaction or epistasis was tested using the R/qtl ‘scantwo’ function (Prins et al. 2010).

When a QTL is found in more than one cross, a joint or a meta-analysis can increase the precision of the QTL location. A joint analysis is considering several populations as one single population, and thus necessitates (1) harmonizing the genotypic and phenotypic data among the populations, then (2) merging the genotypic and phenotypic data into joint matrices, and (3) running the QTL analysis on the joint dataset. On the contrary, in the meta-analysis, a QTL analysis is firstly performed in each population separately, then the resulting scores are combined using the Fisher’s method (Fisher 1932).

To harmonize the genotypic data, as different markers segregated in the four populations, we created a genotype matrix from the union of the four individual data sets. Markers with no data in some of the populations were imputed using the R/qtl ‘Argmax’ function. For joint analysis, only the populations that showed a QTL in single-population analyses were pooled. Meta-analysis was always ran using the four populations pooled.

### The phenotypic data were harmonized using centralized and normalized phenotypic data, that is

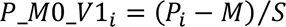

Where *P_i_* is the phenotypic value of the individual *i*, *M* is the average value in the population, and *S* is the standard deviation in the population. Centralized and normalized data always have a mean equal to 0 and a variance equal to 1, making them more suitable for combination in joint analyses.

Both joint-and meta-QTL analyses were run using MapDisto 2.0.

To identify candidate genes in the fine-mapped regions, the MSU Rice genome annotation database and the Overview of Functionally Characterized Genes in Rice Online database (OGRO) (Yamamoto et al. 2012) were used. A 1-LOD drop-off support interval was used to define the search region for each QTL.

## Results

### SNP markers and genetic maps

A total of 332 SNPs were identified as segregating in at least one of the four crosses. Table S1 summarizes the main characteristics of each SNP: position on the genome, allele in the RefSeq and the susceptible and resistant parents, and 250 bp flanking sequences (left and right).

Table 1 summarizes the main statistics of the individual genetic maps obtained for each cross. Overall, we obtained genetic map sizes coherent with their expected size according to ten high-quality maps based on flexible and scalable genotyping by sequencing (fsGBS) (Heffelfinger et al. 2014; Fragoso et al. 2017), the absence of pollen contamination in the self-pollination process and the high quality of the SNP data.

**Table 1.** Summary of genetic maps characteristics in the four F_2_ populations. Map sizes are expressed in centimorgans (cM) calculated with the Kosambi mapping function

| *Cross* | *Population size* | *Number SNPs* | *Number SSRs* | *Total markers* | *Map size (cM)* |
| --- | --- | --- | --- | --- | --- |
| WAS 208 × BBT 50 | 104 | 207 | 2 | 209 | 1,296 |
| Badka × BBT 50 | 105 | 226 | 1 | 227 | 1,570 |
| PTB 25 × BBT 50 | 108 | 184 | 1 | 185 | 1,617 |
| FD 2000 × BBT 50 | 105 | 217 | 0 | 217 | 1,630 |

### Segregation distortion (SD)

SD was observed on chromosome 3 (11.1-21.0 Mbp) in the crosses involving WAS 208, Badka and FD 2000. This region is commonly affected by SD in *indica* × *japonica* crosses (Wang et al. 2009; Fragoso et al. 2017). The homozygotes for the donor allele (AA genotype) were favored over both the homozygotes for the susceptible parent allele (BB genotype) and the heterozygotes (AB) (Figure S1). In the cross WAS 208 × BBT 50 cross, a pattern of SD was also observed on chromosome 8 (3.53-16.44 Mbp), where the AB genotypes were favored over the AA. In the cross PTB 25 × BBT 50, segregation distortion was observed on chromosomes 2 (20.1 Mbp, 47AA:50AB:11BB), 6 (2.9-7.9 Mbp, 34AA:64AB:10BB) and 9 (10.8-12.2 Mbp, 42AA:54AB:12BB).

### RHB incidence and severity variation

In the cross WAS 208 × BBT 50, the parental lines showed 24.2±11.9% and 98.1±3.2% of RHB incidence and 25.8±32.7% and 69.5±37.7% of RHB severity in WAS 208 and BBT 50, respectively. The F_3_ families (each representing one F_2_ parental plant) exhibited a continuous variation of incidence ranged between 21.4% and 100%, as well as a variation of severity ranged between 9% and 68% (Figure 2). The high incidence score for the susceptible parent BBT 50 – almost 100% – and the maximum values in the F_3_ families indicate a perfect infection efficiency. The similarity between the severity score for BBT 50 and the maximum scores in F_3_ families indicates that the experimental design, and in particular the population size, were adequate to capture the range of variation between the parental values.

**Figure 2.**
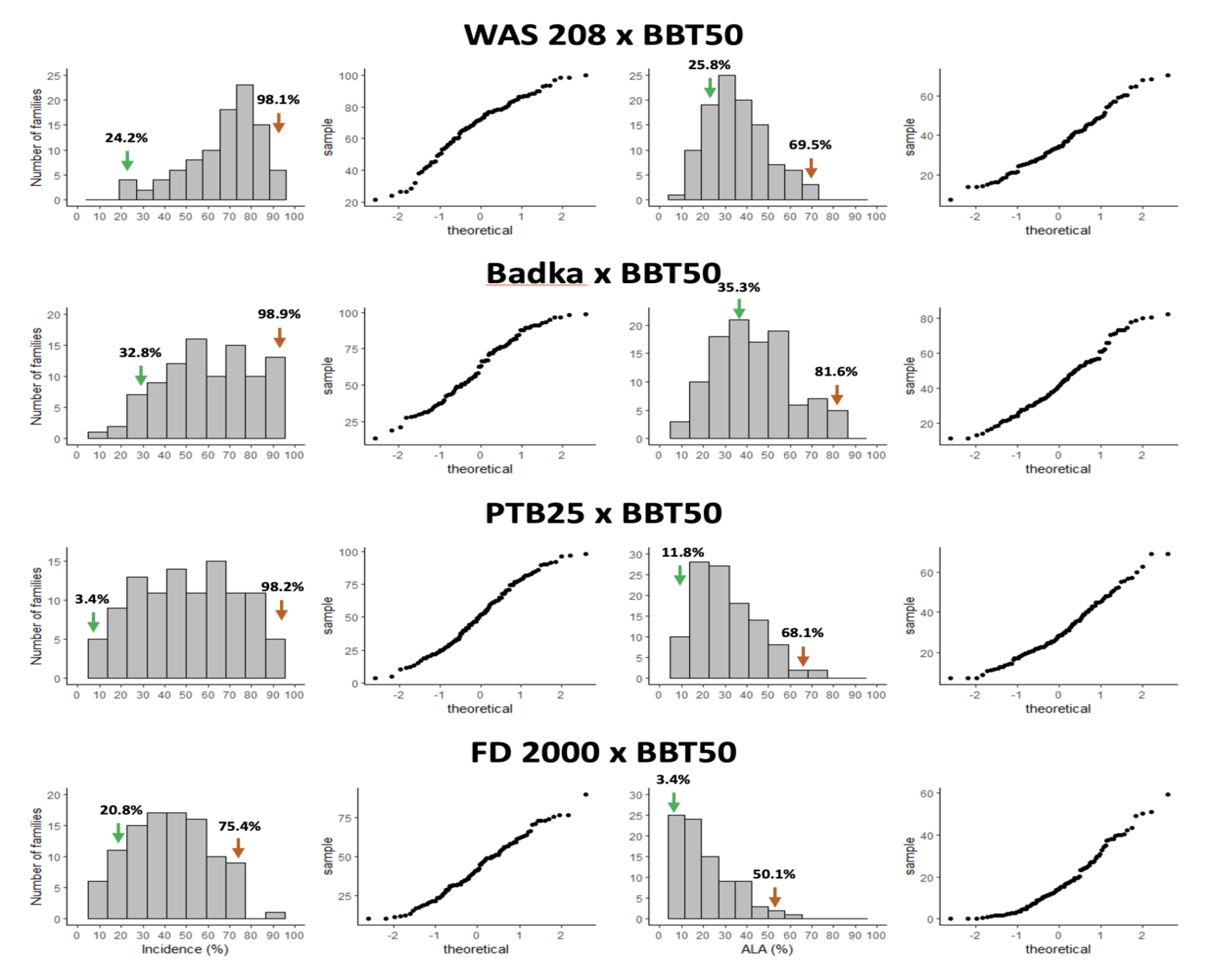
Distribution of the means of RHB incidence (% of plants with symptoms) and severity (% of affected leaf area, ALA) in F_3_ families, in four *indica* × *japonica* rice mapping populations. Associated quantile-quantile plot to the right of every distribution. Means in F_3_ families were calculated from 60 plants (incidence) or 10 plants (ALA). The green and orange arrows represent the average of the resistant and susceptible parents, respectively.

**Figure 3.**
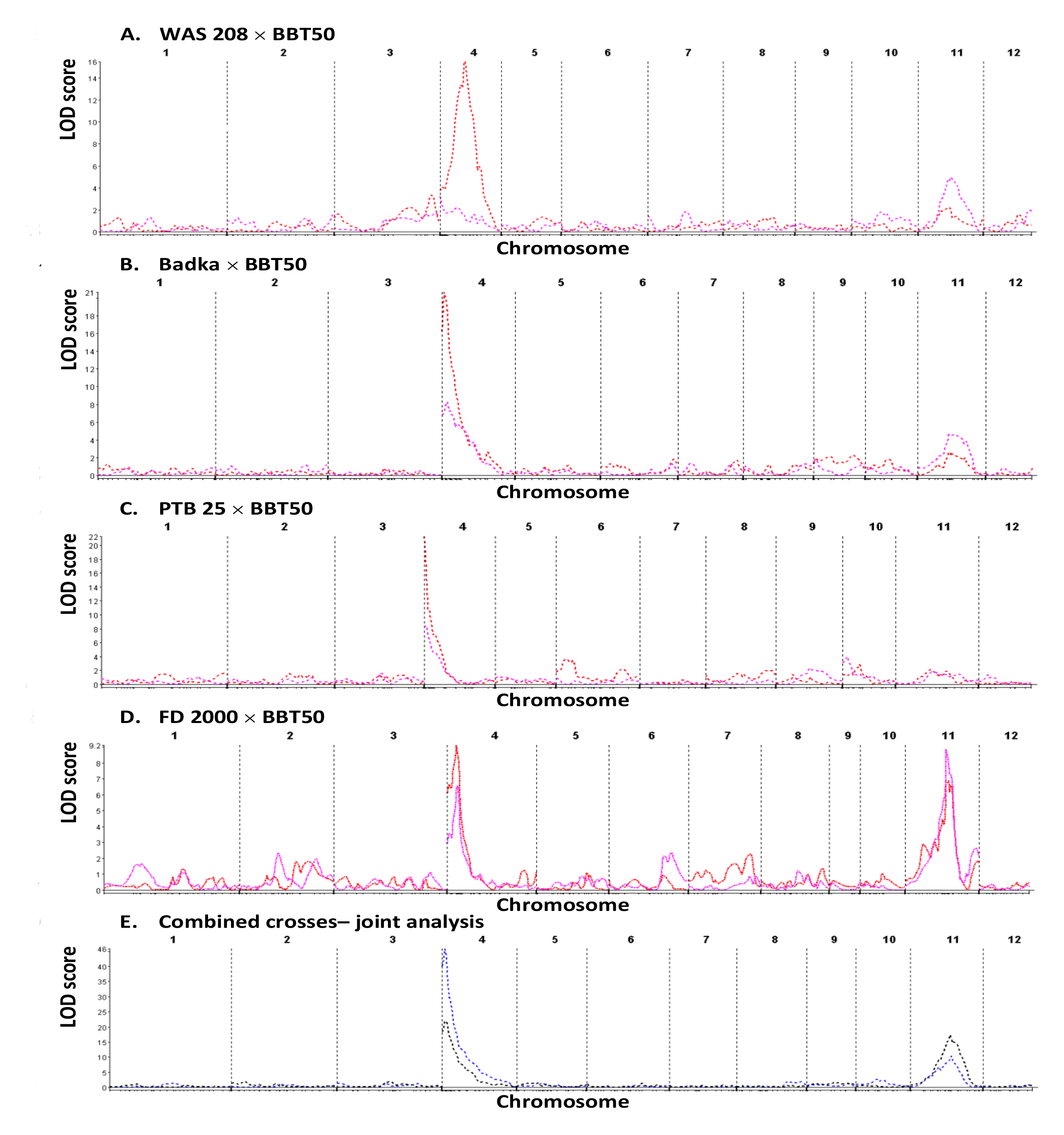
Whole-genome QTL plots (SIM) in the four crosses analyzed independently (A-D), and jointly (E). Red:incidence; purple: severity (ALA)

**Figure 4.**
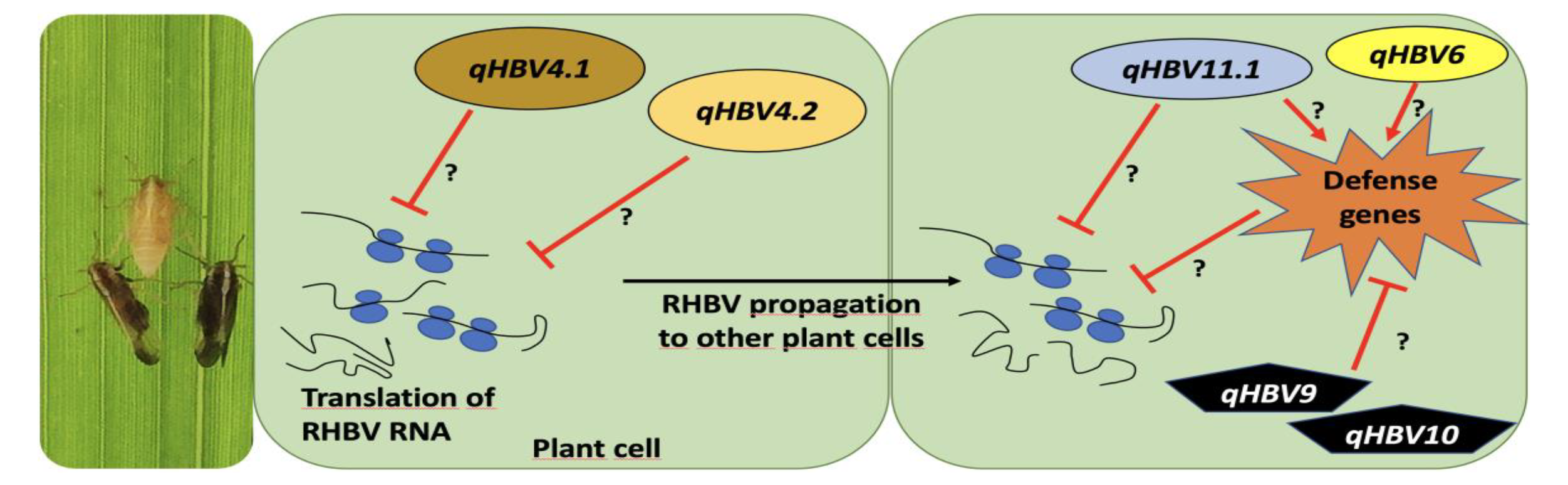
Hypothetical model of plant resistance mechanisms to RHBV

In the cross Badka × BBT 50, the resistant and susceptible parents exhibited 32.8±12.4% and 98.9±2.9% of RHB incidence, respectively. In the same order, these parents displayed 35.3±17% and 81.6±27.7% of severity. The incidence in the F_3_ families ranged between 13% to 98%, and the severity varied between 11% and 80% (Figure 2). A positive correlation between incidence and severity of the disease was detected (*r*=0.56, *p*<0.0001) using the entire population, suggesting a common partial genetic control for both assessments of RHB disease. Some F_3_ families exhibited lower incidence and severity than the resistant parent Badka.

In the cross PTB 25 x BBT 50, the incidence of the resistant and susceptible parents was 3.4±3.5% and 98.2±3.3%, respectively, and ranged between 5% to 96% in the F_3_ families. Furthermore, PTB 25 and BBT 50 showed 11.8±22.1% and 68.1±25.1% of severity, respectively. The same trait varied between 7% and 69% in the F_3_ families.

In the cross FD 2000 × BBT 50, the resistant and susceptible parents showed 20.8±8.8% and 75.4±14.1% of RHB incidence, and 3.4±6.5% and 50.1±33.1% of severity, respectively. In the F_3_ families; incidence ranged between 10% and 90%, and severity between 0% and 59%. The incidence and severity for the susceptible parent BBT 50 were notably lower than in the crosses before mentioned, indicating a low infection efficiency that can affect the power of QTL detection and the QTL effects estimation. In contrast, the incidence of FD 2000 was higher than in previous works (Romero et al. 2014; Cruz-Gallego et al. 2018).

Incidence and severity showed correlation between 0.42 and 0.66 (Table 2), indicating either a common genetic control of the two traits, or that they are interdependent.

**Table 2.**
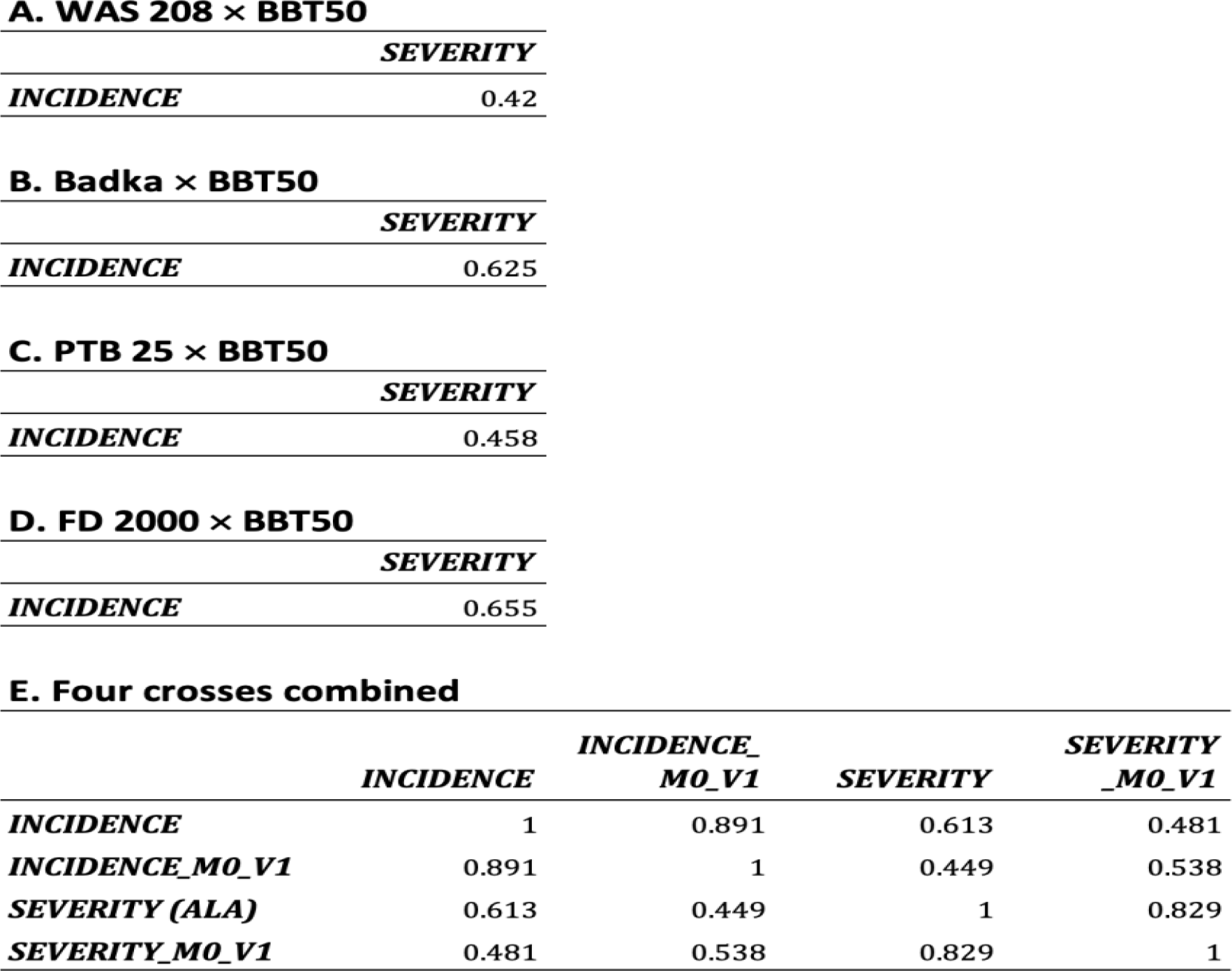
Trait correlation values in each population, and in the four F_2_ populations combined. The '_M0' and '_V1' suffixes stand for normalized and centralized data, respectively

### The major QTL *qHBV4.1* for RHBV incidence is present in most donors

A major QTL, *qHBV4.1,* on chromosome 4 for RHBV incidence was identified by SMR, SIM and CIM in the crosses Badka × BBT 50 (LOD=20.97, R^2^=0.63), PTB 25 × BBT 50 (LOD=21.26, R^2^=0.60) and FD 2000 × BBT 50 (LOD=9.11, R^2^=0.34) (Table 3). It was not detected in the cross WAS 208 × BBT 50, although a fake QTL analysis (Lorieux 2018) showed that it could be present in WAS 208, but with a much smaller effect (data not shown). The QTL support intervals are overlapping in the three populations, suggesting that the same QTL is shared by the three resistance donors. Joint analysis gave a LOD=47.00 and R^2^=0.42 at the position ∼3.56 Mbp. The *qHBV4.1* position also corresponds to a previously identified locus characterized as the major contributor to RHBV resistance in FD 2000 and FD 50 (Romero *et al*., 2014), confirming the wide range of action of this QTL. It explained 34-63% of the trait variance, indicating that *qHBV4.1* is a major regulating factor of incidence of RHBV infection. The estimation of QTL effects for *qHBV4.1* showed that this QTL is mostly of the additive type. The same genomic region was also associated with RHB severity in the same crosses, however with lower LOD scores and R^2^ values (LOD=6.69-8.50, R^2^=25-31%).

**Table 3.**
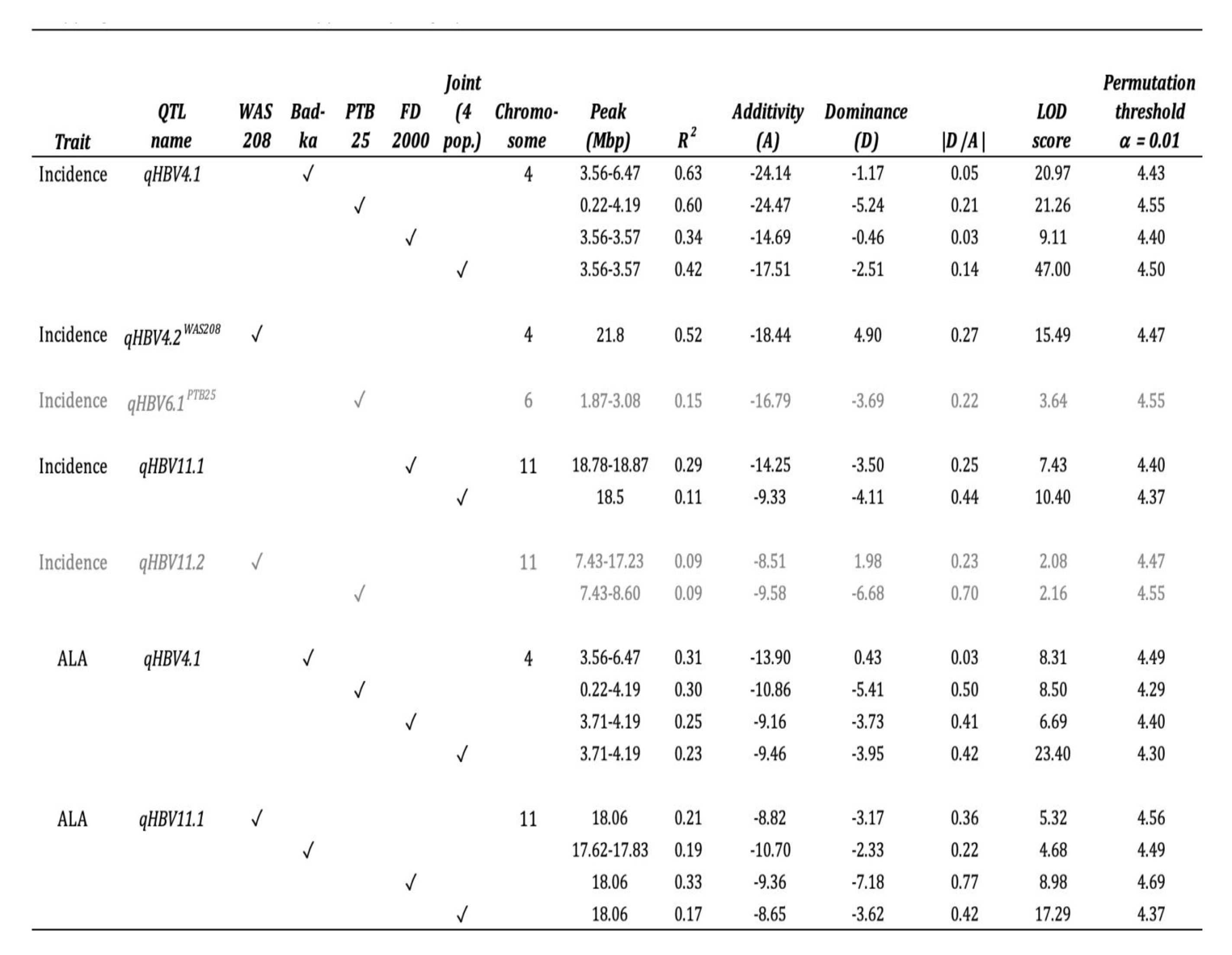
Summary statistics for QTLs detected with Simple Interval Mapping (SIM) analysis in the four F_2_ crosses, and joint analysis, with Qgene version 4.40. Positions in Mbp on Nipponbare genome (MSU v7). Empirical threshold α_0.01_ calculated with 10,000 permutations. R^2^: part of the trait variance explained by the QTL. A: Additivity effect. D: Dominance effect. QTLs that were detected with Composite Interval Mapping (CIM) but not with SIM appear in pale gray

### A new major QTL for RHB incidence, *qHBV4.2*, identified in WAS 208

The control of RHB incidence in WAS 208 was mainly explained by a different QTL on chromosome 4, designated as *qHBV4.2^WAS208^* (LOD=15.49, R^2^=0.52) by SMR, SIM and CIM. This QTL was not found in the other crosses (Table 3). This newly discovered QTL was located between 21.29 and 21.81 Mbp and explained 52% of the incidence variance. *qHBV4.2* is therefore another major QTL for RHB incidence that seems less frequent than *qHBV4.1* in the rice germplasm.

### Two new QTLs for RHB incidence identified in WAS 208 and PTB 25

Two additional QTLs, although of lesser effect, were detected for RHB incidence (Table 4): – *qHBV6.1^PTB25^* on chromosome 6 (0.18-1.76 Mbp, LOD_SIM_=3.64, LOD_CIM_=9.71, R^2^=25%), mostly of the additive type effect and detected in the PTB 25 × BBT 50 cross only. – *qHBV11.2* on chromosome 11 in the crosses involving WAS 208 (7.43-11.9 Mbp, LOD_CIM_=5.02, R^2^=21%) and PTB 25 (7.43-16.6 Mbp, LOD_CIM_=5.9, R^2^=24.2%). However, the SMR or SIM methods produced LOD scores under the retained threshold (Tables 3 and 5).

**Table 4.**
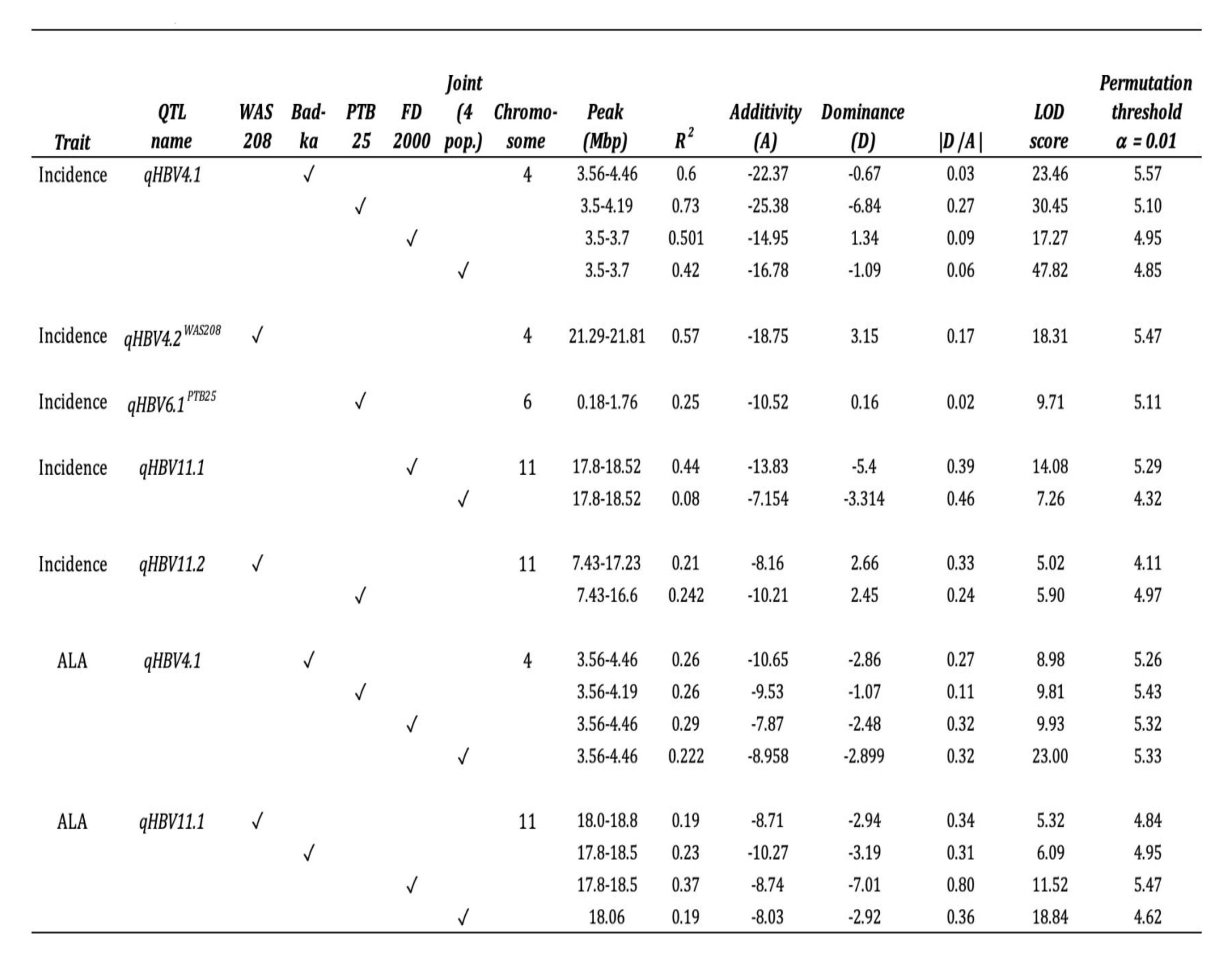
Summary statistics for QTLs detected with Composite Interval Mapping (CIM) analysis in the four F_2_ crosses, and joint analysis, with Qgene version 4.40. Cofactor selection method: Forward selection, automatic, QTL cofactors instead of markers = no. Positions in Mbp on Nipponbare genome (MSU v7). Empirical threshold α_0.01_ calculated with 10,000 permutations. R^2^: part of the trait variance explained by the QTL. A: Additivity effect. D: Dominance effect.

**Table 5.** Summary statistics for QTLs detected with Simple Marker Regression (SMR) analysis in the four F_2_ crosses, joint- and meta-analyses, with MapDisto version 2.0. Positions in Mbp on Nipponbare genome (MSU v7). Empirical threshold α_0.01_ calculated with Qgene 4.40, with 10,000 permutations*. R^2^: part of the trait variance explained by the QTL. A: Additivity effect. D: Dominance effect. QTLs that were detected with Composite Interval Mapping (CIM) but not with SMR appear in pale gray

| Trait | QTL name | WAS 208 | Bad-ka | PTB 25 | FD 2000 | Joint (4 pop.) | Meta (4 pop.) | Chromosome | Peak (Mbp) | R<sup>2</sup> | Additivity (A) | Dominance (D) | \|D / A\| | F-test | -Log<sub>10</sub> (p-value of test) | Permutation threshold $\alpha = 0.01$ |
| --- | --- | --- | --- | --- | --- | --- | --- | --- | --- | --- | --- | --- | --- | --- | --- | --- |
| Incidence | qHBV4.1 |  | ✓ |  |  |  |  | 4 | 3.56-6.47 | 0.61 | 22.43 | -1.65 | 0.07 | 74.18 | 19.33 | 4.35 |
|  |  |  |  | ✓ |  |  |  |  | 0.22-4.19 | 0.60 | 24.47 | -5.24 | 0.21 | 77.85 | 20.67 | 4.18 |
|  |  |  |  |  | ✓ |  |  |  | 3.56-3.57 | 0.34 | 14.69 | -0.46 | 0.03 | 25.29 | 8.84 | 4.20 |
|  |  |  |  |  |  | ✓ |  |  | 3.56-3.57 | 0.42 | 17.51 | -2.51 | 0.14 | 142.30 | 46.65 | 4.29 |
|  |  |  |  |  |  |  | ✓ |  | 3.56-3.57 | - | - | - | - | - | 47.00 | * |
| Incidence | qHBV4.2<sup>WAS208</sup> | ✓ |  |  |  |  |  | 4 | 21.8 | 0.50 | 17.67 | 4.40 | 0.25 | 48.04 | 14.42 | 4.28 |
| Incidence | qHBV6.1<sup>PTB25</sup> |  |  | ✓ |  |  |  | 6 | 2.97-3.08 | 0.13 | 15.04 | -6.76 | 0.45 | 7.82 | 3.16 | 4.31 |
| Incidence | qHBV11.1 |  |  |  | ✓ |  |  | 11 | 18.78-18.87 | 0.29 | 14.09 | -2.92 | 0.21 | 19.45 | 7.10 | 4.22 |
|  |  |  |  |  |  | ✓ |  |  | 18.52-18.87 | 0.11 | 9.33 | -4.11 | 0.44 | 25.24 | 10.32 | 4.29 |
|  |  |  |  |  |  |  | ✓ |  | 18.52-18.87 | - | - | - | - | - | 8.91 | * |
| Incidence | qHBV11.2 | ✓ |  |  |  |  |  | 11 | 7.43-17.23 | 0.10 | 7.55 | 2.05 | 0.27 | 5.12 | 2.11 | 4.29 |
|  |  |  |  | ✓ |  |  |  |  | 7.43-8.60 | 0.09 | 9.15 | -6.09 | 0.67 | 5.03 | 2.08 | 4.25 |
| ALA | qHBV4.1 |  | ✓ |  |  |  |  | 4 | 3.56-6.47 | 0.29 | 12.85 | -1.09 | 0.09 | 21.12 | 7.67 | 4.23 |
|  |  |  |  | ✓ |  |  |  |  | 0.22-4.19 | 0.30 | 10.54 | -2.48 | 0.24 | 22.09 | 8.01 | 4.18 |
|  |  |  |  |  | ✓ |  |  |  | 3.71-4.19 | 0.25 | 8.86 | -3.19 | 0.36 | 17.23 | 6.45 | 4.38 |
|  |  |  |  |  |  | ✓ |  |  | 3.56-3.71 | 0.23 | 9.46 | -3.95 | 0.42 | 60.94 | 23.23 | 4.22 |
|  |  |  |  |  |  |  | ✓ |  | 3.70-4.19 | - | - | - | - | - | 20.75 | * |
| ALA | qHBV11.1 | ✓ |  |  |  |  |  | 11 | 18.06 | 0.21 | 8.82 | -3.17 | 0.36 | 13.43 | 5.17 | 4.31 |
|  |  |  | ✓ |  |  |  |  |  | 17.62-17.83 | 0.19 | 10.70 | -2.33 | 0.22 | 11.62 | 4.55 | 4.16 |
|  |  |  |  |  | ✓ |  |  |  | 18.06 | 0.32 | 9.22 | -6.66 | 0.72 | 23.72 | 8.46 | 4.38 |
|  |  |  |  |  |  | ✓ |  |  | 18.06 | 0.17 | 8.65 | -3.62 | 0.42 | 43.49 | 17.16 | 4.30 |
|  |  |  |  |  |  |  | ✓ |  | 18.06 | - | - | - | - | - | 15.40 | * |

\*Meta-analysis not available in Qgene

### A new QTL, *qHBV11.1*, controls RHB severity

One of the most interesting results of this work is the discovery of a new QTL associated with RHB symptoms severity (measured as ALA). It was detected on chromosome 11 in the crosses WAS 208 × BBT 50 (18.0-18.8 Mbp, LOD=5.32, R^2^=0.21), Badka × BBT 50 (17.8-18.5 Mbp, LOD=4.68, R^2^=0.19) and FD 2000 × BBT 50 (17.8-18.5 Mbp, LOD=8.98, R^2^=0.33). This QTL was designated as *qHBV11.1* and explained 17-33% (SIM) or 19-37% (CIM) of the trait variation, depending on the cross (Tables 3 and 4). The QTL effects in the populations involving WAS 208 and Badka indicate an additive behavior of *qHBV11.1*, while in FD 2000 × BBT 50 it seems to be more dominant (Table 3). In the cross FD 2000 × BBT 50, *qHBV11.1* was also significant for RHB incidence, although with lower statistics (LOD=7.43, R^2^=0.29). Interestingly, *qHBV11.1* was significant for ALA_strict (LOD=4.05), which suggests that two distinct genes act onto RHB severity, possibly in interaction.

### **The *qHBV4.1* and *qHBV11.1* QTLs show strong interaction**

Testing interaction between QTL regions with the R/qtl “scantwo” function produced a strong signal between the two QTLs *qHBV4.1* and *qHBV11.1* (LOD>15) (see example for the FD 2000 × BBT 50 cross in Figure S2), revealing either an epistasis relationship between the two regions, or a simple interdependency between the two traits. This is coherent with the positive correlation observed between severity and incidence.

### Joint-and meta-analyses provide good candidates for QTL cloning

The joint and meta-analyses approaches allow pooling populations, providing more resolution for QTL mapping. This allowed us identifying candidate genes that could underlie two of the QTLs we discovered.

### The qHBV4.1 region contains the AGO-4 Argonaute gene

In the *qHBV4.1* region, 33 genes encoding proteins with unknown function, five hypothetical genes, four genes encoding different types of kinases, two genes of the MEG (maternally expressed gene) family involved in the translocation of nutrients to the seed, and one *Argonaute* gene were identified. The most interesting gene within the *qHBV4.1* region is probably the *Argonaute* AGO-4 (MSU: LOC_Os04g06770, Chr4:3,562,793-3,555,220bp). AGO proteins are effector proteins of RNA silencing pathways, which regulate gene expression in a sequence-specific manner (Duan et al. 2015). RNA silencing is also the main antiviral defense mechanism possessed by plants. It can be post-transcriptional (PTGS) or transcriptional (TGS) (Carbonell and Carrington 2015).

### The qHBV11.1 region contains a gene for durable resistance to Rice stripe virus

The genomic region of *qHBV11.1* contains many nucleotide binding site-leucine-rich repeat (NBS-LRR) genes, which are broadly known to confer resistance to multiple diseases (McHale et al. 2006). Other types of genes are also found in the interval, and remarkably the *STV11* gene (MSU:LOC_Os11g30910, Chr11:17,984,964-17,986,719b; RAP-DB: Os11g0505300), which confers durable resistance to *Rice stripe virus* (RSV), one of the most devastative viral diseases of rice in Asia (Wang et al. 2014). Interestingly, this virus belongs to the same genus as RHBV and it is also transmitted by planthoppers (*Laodelphax striatellus* Fallen). It should be also noted that, very close to the *qHBV11.1* support interval, there are several paralog histidine kinase/Hsp90-like ATPase genes (MSU:LOC_Os11g31480, MSU:LOC_Os11g31500) that also confer resistance to RSV (Hayano-Saito and Hayashi, 2020).

## Discussion

Successful genotype-phenotype association studies require precise methodologies to assess phenotypic variables. The latest technological advances in image acquisition have allowed the development of computer tools for picture processing and analysis that provide accurate data. Image processing for quantifying damage caused by plant diseases is particularly useful as it advantageously replaces traditional scoring scales, which are too dependent on the observer. The RHB disease has been commonly evaluated using the percentage of diseased plants at a given time – that is, the incidence. However, the main drawback of this approach is that it classifies all the plants with symptoms in the same class, without considering the level of damage – the severity. Understanding the genetic mechanisms of resistance to the *hoja blanca* disease therefore requires proper assessment of not only its incidence but also its severity. In this sense, we proposed for the first time the use of digital images to evaluate the severity of damage caused by RHB by estimating the affected leaf area. We showed that severity varies continuously among F_3_ families, a characteristic behavior of quantitative features. Severity and incidence evaluation allowed us to identify different QTLs, allowing us to better explain the complexity of the genetics of resistance to RHB. Keeping in mind that the parents of the four populations were among the most resistant of a larger panel (Cruz et al. 2018), and although significant differences between incidence and severity were found among the parents of the four populations, complete resistance was not observed, confirming a previous observation by (Morales and Jennings 2010) about the lack of complete resistance to RHB in the *O. sativa* genepool.

### RHB resistance is regulated by multiple QTLs

The QTLs found in this study show that resistance to RHB, assessed both by affected leaf area and incidence, is controlled by multiple genes. The co-location of LOD score peaks for incidence in the same region of chromosome 4 in three different crosses suggests that it is likely the same QTL, namely *qHBV4.1*. Although *qHBV4.1* was associated to both phenotypic variables, its association was greater with incidence where it explained up to 63.6% of the phenotypic variance. These results suggest that the main action of *qHBV4.1* could take place in the first phase of the interaction of the virus with the plant, preventing the virus to enter and propagate in the plant, which is reflected in a smaller number of plants with symptoms. In addition, the little variation in nucleotides – less than 1% – in the region of *qHBV4.1*, between Badka and PTB 25 indicates a probable common local ancestry (Figure S3). The differences in the LOD score (5.83 *vs.* 8.26) and in R^2^ values (21.4% *vs.* 36.6%) for ALA, in these two populations, could be explained by the difference in the percentage of virulence of the vector colonies, which were ∼69% and ∼46% for Badka × BBT 50 and PTB 25 × BBT 50, respectively. Also, the phenotypic evaluation trials were performed at different times of the year for the different populations. This involves the use of different vector colonies, and implies variations in microenvironmental factors such as day and night temperature, radiation or humidity that might affect the behavior of the insect and its ability to transmit the virus.

A major QTL had been previously identified in the region of *qHBV4.1* in the varieties FD 2000 and Fedearroz 50 by incidence assessment (Romero et al. 2014), which suggests a predominant, common mechanism of resistance to RHBV in the different clades of *Oryza sativa*. The genetic variation between the parents observed in this region shows that they carry different alleles, of *japonica* and *indica* origins. This has implications for crop improvement, allowing breeders to broaden the genetic base of resistance in elite germplasm. A QTL for resistance to *Rice stripe virus* (RSV) has been identified between 4.4 and 6.9 Mpb in the N22 Aus variety, explaining 13.4% of the trait variance (Wang et al. 2013). The *qHBV4.1* QTL region might thus contain at least two tenuivirus resistance genes. As mentioned above, in the region of *qHBV4.1* there is a putative candidate gene that encodes for the AGO-4 Argonaute protein (LOC_Os04g06770). AGO proteins, in addition to be regulatory factors of endogenous gene expression, also play a critical role in the defense against viruses through interference with small RNA of viral origin which bind to AGOs and serve as a guide for it to cut new viral RNA particles (Mallory and Vaucheret 2011)(Silva-Martins et al., 2020). This system is a common defense mechanism against pathogens, and AGO-4 might well be associated with resistance to RHBV. Moreover, Bhattacharjee et al. (2009) showed, in *Nicotiana benthamiana*, that Argonaute proteins can interact with viral transcripts to alter virus resistance mediated by proteins containing nucleotide-binding/leucine-rich repeats domains. We are currently carrying CRISPR-Cas9 knock-out experiments in AGO-4 in order to test this hypothesis.

A distinct QTL for RHB incidence on chromosome 4, *qHBV4.2^WAS^*, was identified in the WAS 208 × BBT 50 cross. This QTL was also a major one, explaining ∼50% of the phenotypic variation. The region contains genes associated with cold tolerance (SAPK7) (Basu and Roychoudhury, 2014) and photo-oxidative stress (OsAPX7)(Caverzan et al. 2014), and a more interesting gene encoding for a zinc finger-type C3H protein (LOC_Os04g35800). This class of proteins has been found to be associated with resistance viruses in animals through degradation of viral RNA (Gao et al. 2002; Mao et al. 2013). Although there are no reports so far of its involvement in resistance to viruses in plants, it has been found to be associated with response to other types of pathogens (and abiotic stress) (Deng et al. 2012; Jan et al. 2013), which makes it an interesting candidate for RHB resistance. Another candidate is a gene encoding for a MAPKKK kinase, which belongs to a signaling cascade that plays a major role in disease resistance in eukaryotes. This gene is also an interesting candidate because the activation of MAPK proteins occurs in the earliest stages after the interaction of the pathogen with the plant (Meng and Zhang 2013), which coincides well with the identification of this QTL through the assessment of incidence, but not severity, indicating that it – like *qHBV4.1 –* could exert its action in the early stages of the interaction of RHBV with the plant.

In contrast with the two QTLs identified on chromosome 4 that are predominantly associated with incidence, *qHBV11.1* has a greater effect on disease severity. This could suggest a later action, in a second stage of the interaction of the plant with the virus or the insect vector, after the virus manages to enter the cells and to propagate. In this scenario *qHBV11.1* would have a key role in diminishing the damage caused by the virus. This is thus more a “tolerance QTL”.

It has been proposed that the resistance to RHB comes mostly from *japonica* germplasm (Morales and Jennings 2010). If this was the case, and to explain that the resistance sources studied here are of *indica* type, one would need to assume that the region of *qHBV11.1* was inherited from *japonica* through introgression into *indica*. This is not supported by the clustering analysis, which showed that the WAS 208 and Badka parents, for this region, are grouped together with *indica* type accessions such as IR8. Therefore, this QTL does not originate from the *japonica* cluster.

A search for candidate genes for *qHBV11.1* evidenced a co-localization with a QTL for resistance to *Rice stripe virus* (RSV) found in different genetic backgrounds: in the *indica* varieties Teqing (*qSTV11^TQ^*) (Wu et al. 2011) and Shingwang (*qSTV11^SG^*) (Kwon et al. 2012), the Aus varieties Kasalath (*qSTV11^KAS^*) (Zhang et al. 2011), N22 (*qSTV11.1*) (Wang et al. 2013) and Dular (Wu et al. 2011), the *japonica* Kanto 72 (Maeda et al. 2006), and *O. rufipogon* (Wang et al. 2014). Whether those QTLs correspond to the same locus or to different, linked genes still needs to be clarified. The hypothesis of a common defense mechanism acting against RHBV and RSV seems plausible, as both are RNA viruses of the same genus, although it has been found that RHBV is more related to other tenuiviruses like *Echinochloa hoja blanca virus* (EHBV) and *Urochloa hoja blanca virus* (UHBV) than to RSV (Fauquet et al., 2005), besides not being serologically related (Morales and Jennings 2010). Further studies of fine mapping and cloning of *qSTV11^KAS^* showed that the gene responsible for resistance to RSV encodes a sulfotransferase that catalyzes the conversion of salicylic acid to its sulfated form. Although it is not clear how this process confers resistance against the virus, it was found that the susceptible allele of this gene is not capable of inducing this conversion (Wang et al. 2014). Salicylic acid has been found to be essential in the initiating signal to activate systemic resistance against *Tobacco mosaic virus* (Zhu et al. 2014), and also plays a central role in the hypersensitive response against *Potato virus Y* (PVY) (Baebler et al. 2014). It has even been reported as a direct inhibitor of the replication of *Tomato bush dwarf virus* (TBSV) (Tian et al. 2015). It is thus plausible that the *STV11* gene is involved in resistance to RHBV. In all cases, Wang et al demonstrated that the *STV11* gene inhibits the replication of the RSV, which could explain the fact that the QTL LOD score is higher for ALA than for incidence in Badka and WAS208. We are currently analyzing knock-out lines obtained by CRISPR-Cas9 in STV11 to verify this hypothesis.

A second QTL on chromosome 11, *qHBV11.2*, was identified for disease incidence in the two crosses involving WAS 208 and PTB 25. Based on the variation found in the clustering analysis – and assuming the QTL is the same in the two crosses – it is possible that WAS 208 and PTB 25 have different allelic variants of *qHBV11.2* with a different effect on disease resistance, which could explain that in PTB 25 it was also associated with severity. The region could also correspond to two distinct, linked QTLs in the two populations, since their support intervals are quite large (> 3 Mpb). To answer this question, a fine mapping study of the region using larger F_2_ populations would be needed.

An additional QTL for RHB incidence, *qHBV6*, was identified In the PTB 25 × BBT 50 cross on chromosome 6. In a previous GWAS experiment, the same region was identified in the genotypes PTB 25 and Pokkali (Cruz-Gallego et al. 2018), also supported by a biparental QTL analysis in the F2:3 cross FD 50 × WC 366 (our unpublished data). A search for genes in this region found the OsBBI1 gene that participates in various biological processes, among which is the innate immune response. OsBBI1’;s expression is induced by the fungus *Magnaporthe oryzae* and chemical inducers such as salicylic acid (Li et al. 2011), which, as discussed above, is also associated with systemic resistance to viruses. OsBBI1 is thus a good candidate gene for *qHBV6*.

### Resistance QTLs in the susceptible parent

In the PTB 25 × BBT 50 population, two QTLs for RHB incidence were identified on chromosomes 9 and 10, of which the allele that reduces the affected leaf area is brought by BBT 50. One explanation could be that PTB 25 carries susceptibility alleles at these QTLs. However, this is unlikely since inactivation of a susceptibility gene by the resistant allele results in increased resistance. Genetically, the resistant allele of a susceptibility gene is therefore generally recessive. This is not what we observed, since the mean ALA of the heterozygous class AB is more similar to the susceptible class AA (PTB 25) than to the BB class (BBT 50). Therefore, the most likely explanation is that the three other resistant parents (FD 2000, WC 366 and Badka) and the susceptible parent (BBT 50) carry resistant alleles at these QTL, while the PTB 25 parent does not.

### Integration of QTLs into a RHB disease resistance model

Based on the QTLs associated with RHB resistance, we propose a simplistic model that draws the possible processes involving these loci. The QTLs *qHBV4.1* and *qHBV4.2* were found to be most associated with disease incidence, so it is likely that they are involved in the first phase of virus-plant interaction. These QTL may be inhibiting the translation of viral RNA, either by direct action on them or through other genes for degradation. The effect of these QTLs on the incidence, indicates that this mechanism might be the most important for the resistance of the plant to the virus. In the scenario where the favorable alleles of *qHBV4.1* or *qHBV4.2* are not present or that the virus manages to overcome this barrier and to propagate in the plant, an additional mechanism involving *qHBV11.1* would be hampering the multiplication of the virus. This could occur by direct action, inhibiting the synthesis of new viral particles, or more likely, serving as a trigger signal for the activation of defense genes of the systemic resistance system. Although this mechanism would not provide complete resistance to the virus, it would considerably reduce the damage – meaning less leaf area affected. The *qHBV6* QTL is also related to the severity of the disease, so it is possible that it also participates in this same mechanism.

## Conclusion

Altogether these results show that resistance to RHB disease is controlled by multiple quantitative genetic factors of different origins, with varying effects and action mode. The identification of strong candidate genes underlying the detected QTLs supports the idea that resistance is mediated by different defense mechanisms such as viral gene silencing and the salicylic acid pathway. This hypothesis has proven true for other study models such as in the *Citrus tristeza virus* (CTV) where it has been found that silencing of key genes in these defense pathways increases the spread of the virus and its accumulation in the plant (Gómez-Muñoz et al. 2017). Regarding the identification of the genetic mechanisms of resistance, our study lacks long-reads based sequencing data in the resistant and susceptible parental lines. Generating such data would allow structural analysis to identify copy number variations for the candidate genes, as well as comparison of resistant and susceptible alleles at the nucleotide and aminoacid levels.

A considerable amount of work is still needed to understand the fine mechanisms behind defense against RHB. In particular, it will be necessary to clarify whether some of the QTLs found in this study are actually acting against the insect vector, and not the virus itself. Indeed, insect resistance by antixenosis and/or antibiosis are known to be present in rice (see Figure 1 in Fujita et al. 2013) and overlap with some of our QTLs.

Finally, the QTLs detected in this study will benefit the rice breeder’;s community. Marker-assisted selection can be used to combine several resistance QTLs to HBV with other traits, using for instance a pyramiding approach, or other types of marker-assisted selection.

### List of abbreviations

ALA: affected leaf area
Badka: Badkalamkati variety, accession number IRGC 45011
BBT 50: Bluebonnet 50 variety
CIAT: Centro Internacional de Agricultura Tropical (now Alliance Bioversity-CIAT)
CIM: composite interval mapping
cM: centimorgan
CTV: *Citrus tristeza virus*
DAI: days after infestation
DNA: deoxyribonucleic acid
EDTA: ethylenediaminetetraacetic acid
EHBV: *Echinochloa hoja blanca virus*
FD 50: Fedearroz 50 variety, accession number IRGC 1799-1-1
FD 2000: Fedearroz 2000 variety, accession number IRGC 124388
FLAR: Fondo Latinoamericano para Arroz de Riego
fsGBS: flexible and scalable genotyping by sequencing
IFC: integrated fluidic circuit
LOD: logarithm in base 10 of likelihoods odds
MATAB: mixed alkyltri-methylammonium bromide
MEG: maternally expressed gene
MLGM: generalized linear mixed model
NBS-LRR: nucleotide binding site-leucine-rich repeat
PCR: polymerase chain reaction
PTGS: post-transcriptional gene silencing
PTB 25: PTB 25 variety, accession number IRGC 6386
PVY: *Potato virus Y*
QTL: quantitative trait locus
RHB: rice *hoja blanca* disease
RHBV: *Rice hoja blanca virus*
RNA: ribonucleic acid
RSV: *Rice stripe virus*
SD: segregation distortion
SIM: simple interval mapping
SMR: single-marker regression
SNP: single nucleotide polymorphism
TBSV: *Tomato bush dwarf virus*
TGS: transcriptional gene silencing
UHBV: *Urochloa hoja blanca virus*
WAS 208: WAS 208-B-B-5-1-1-3 variety, accession number IRGC 121855
WGS: whole-genome sequencing

## Declarations

### Ethical Approval and Consent to participate

Not applicable

### Consent for publication

Not applicable

### Availability of supporting data

The datasets used and/or analyzed during the current study are available from the corresponding author on reasonable request.

### Authors’; contributions

AS wrote the draft manuscript and performed the SNP design and QTL analysis, MEM set up the disease severity experiments, CQ supervised the SNP genotyping, JQ did the statistical treatment of phenotype data, JT led the Agrobiotechnology laboratory, EG led the FLAR, MC supervised the phenotyping experiments, ML designed the QTL experiments and edited the manuscript, MC and ML supervised AS work.

## Acknowledgements

We warmly thank Drs. Laurence Albar, Sébastien Cunnac, Mathilde Hutin and Marlène Lachaux from the IRD for their useful comments on the manuscript. We also thank the two reviewers for their careful review that helped significantly improving the quality of the manuscript, and the associate editor for handling the submission.

## Supplemental figures

**Figure S1.**
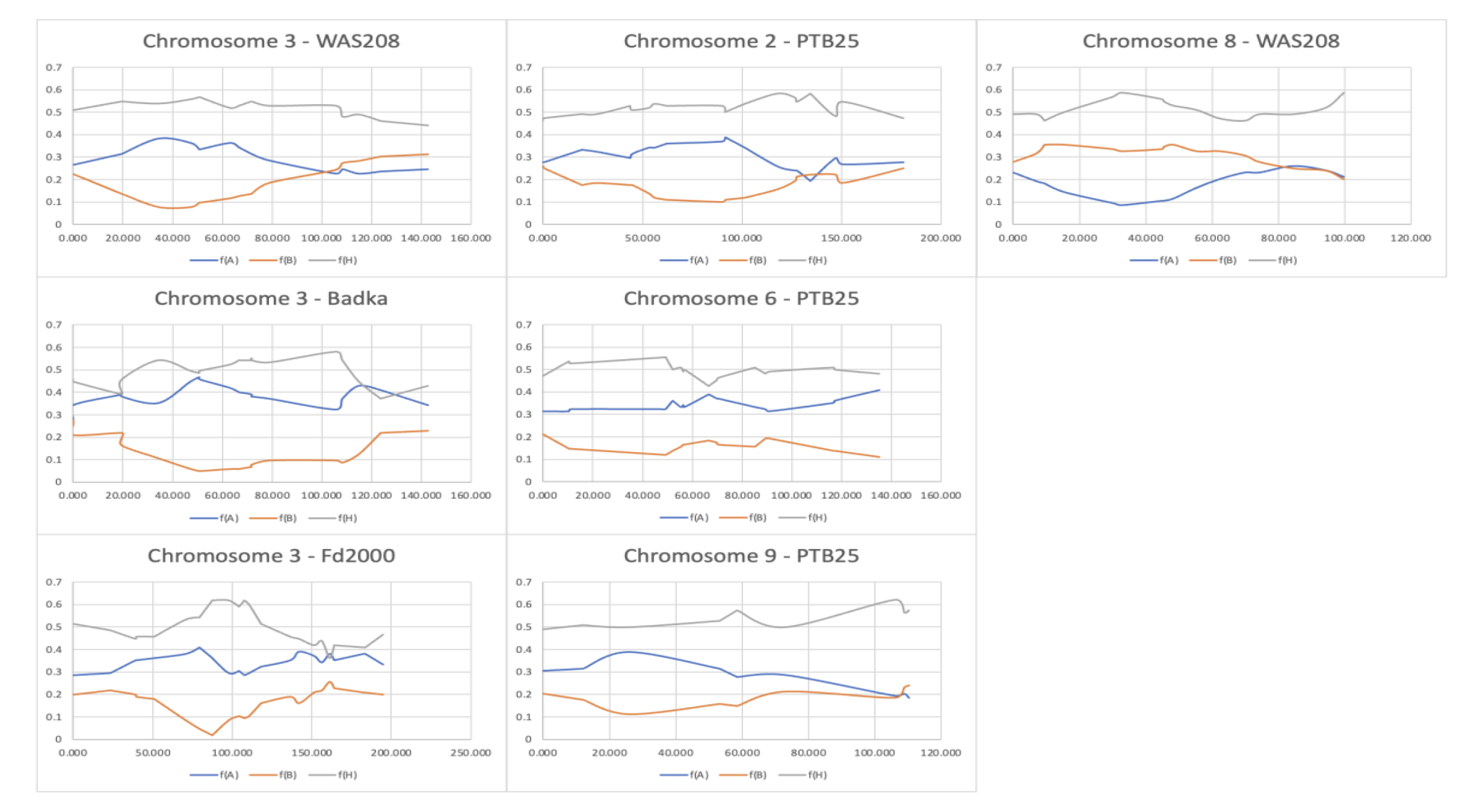
Segregation distortion in several chromosomes, in the F_2_ four populations. The X axis represents the position in cM on the chromosome. The Y axis is the frequency of the three F_2_ genotypes: f(A): homozygous female parent (resistant to HBV); f(B): homozygous male parent BB 50 (susceptible to HBV); f(H): heterozygous. One clearly sees the departure from the expected genotypic frequencies, which in the F_2_ cross are 0.25, 0.25 and 0.5 for f(A), f(B) and f(H), respectively.

**Figure S2.**
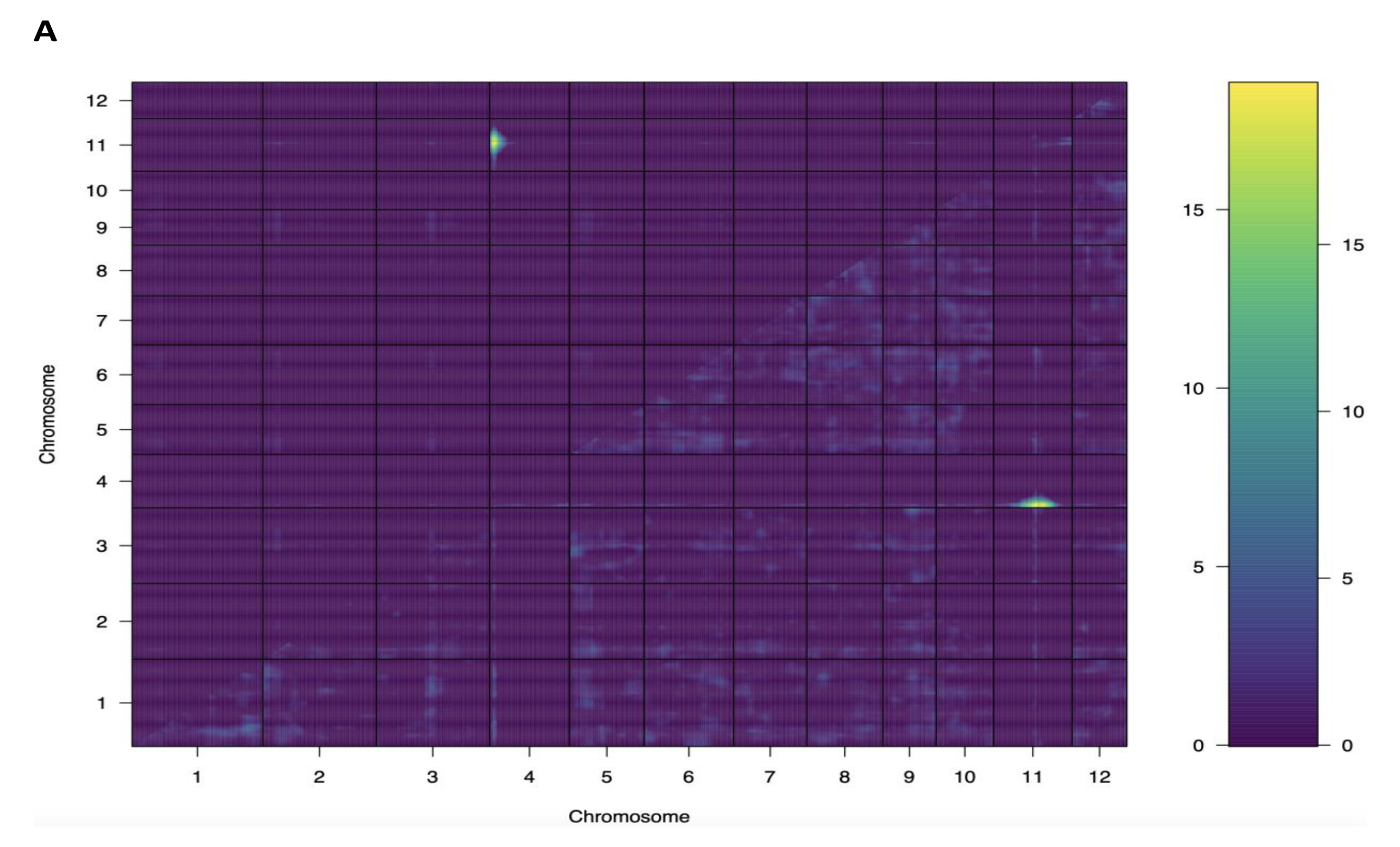
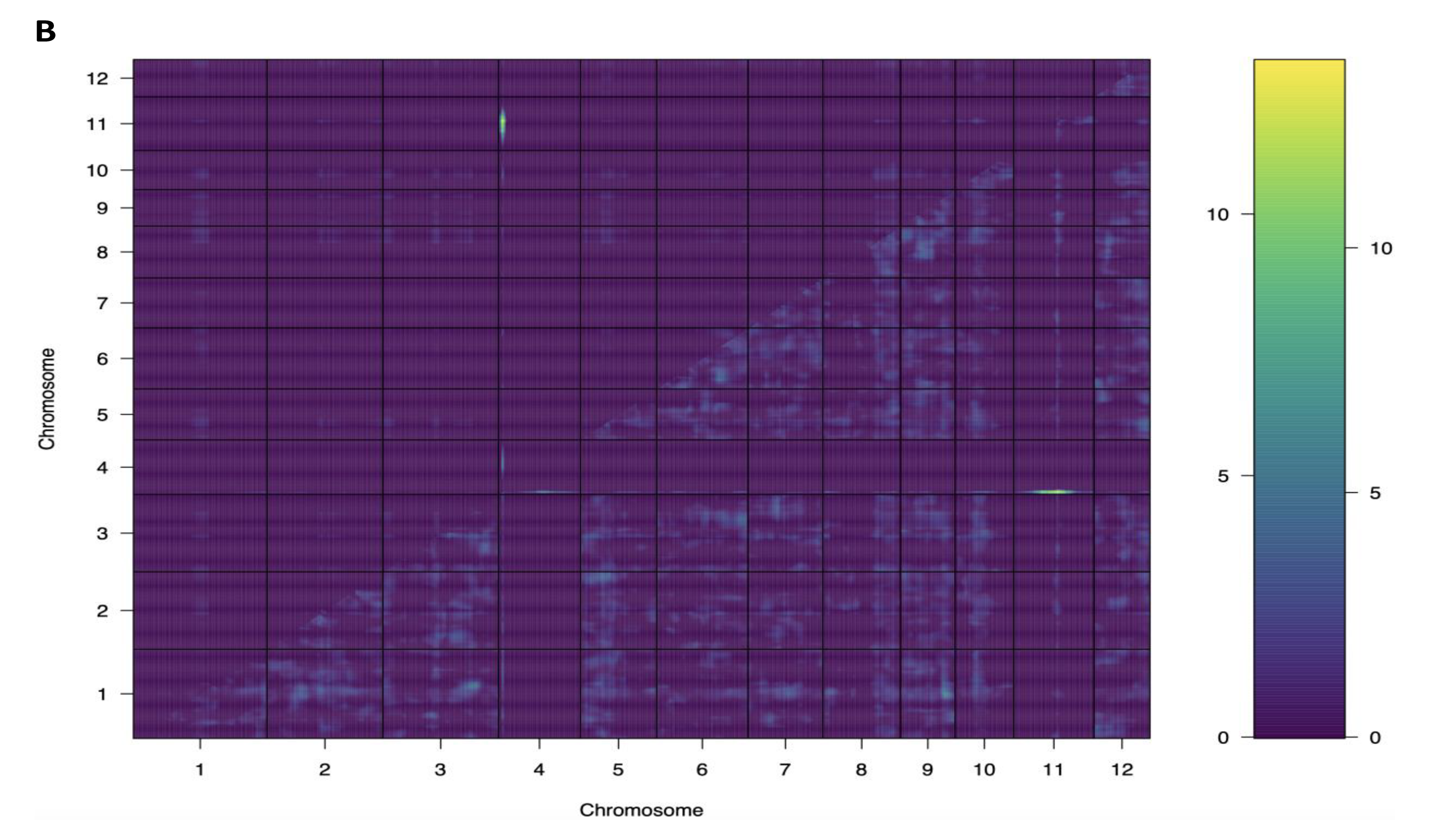
2-D plot for interaction analysis between genomic regions calculated in the FD2000 × BBT50 cross. A: RHBV incidence. B: RHBV severity

**Figure S3.**
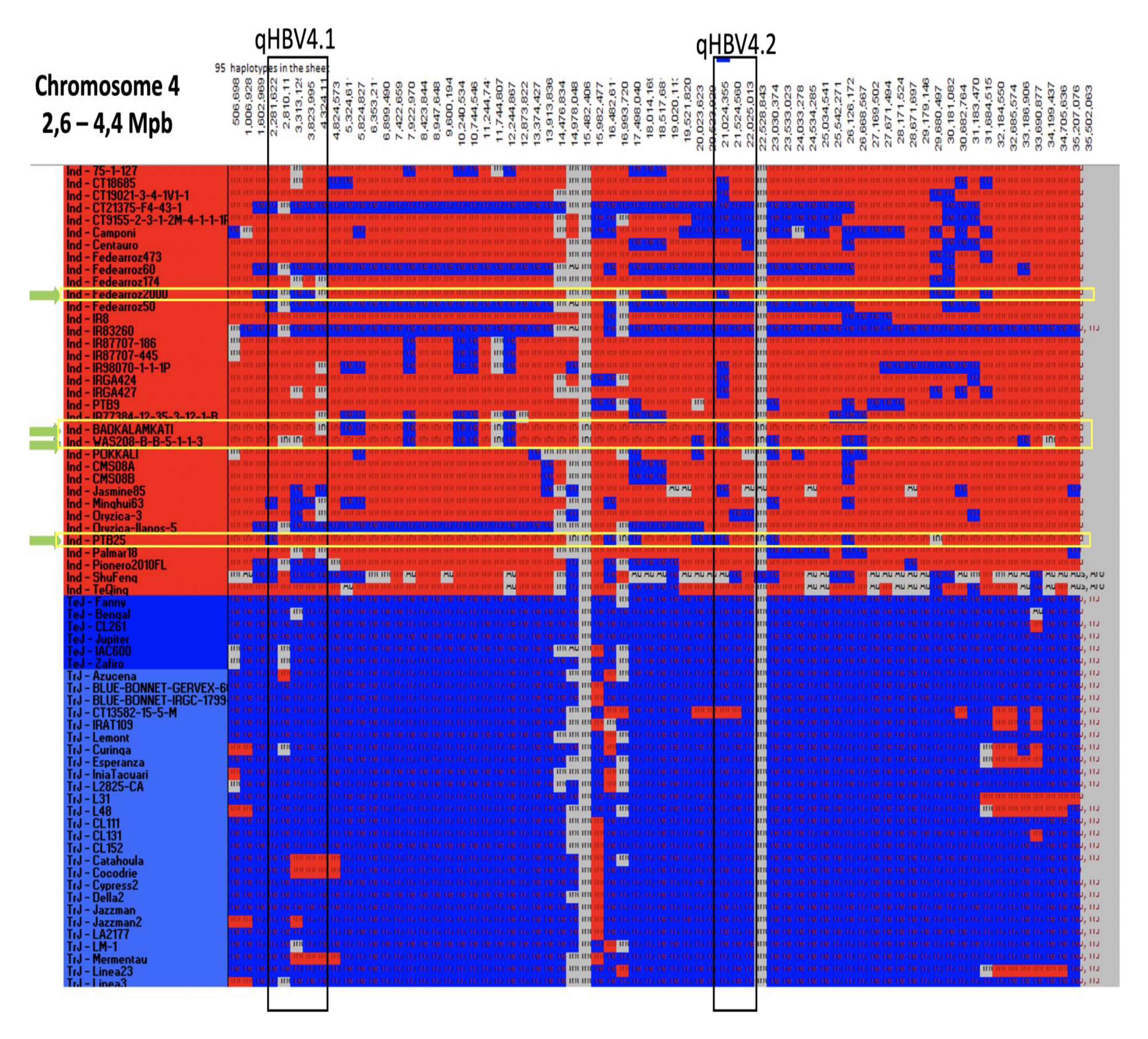
Graphical genotyping showing probable local ancestry for the four RHB-resistant parents (green arrows), in the two QTL regions *qHBV4.1* and *qHBV4.2* on chromosome 4. X-axis: physical coordinates (in bp) along chromosome 4, spanning the region 2.6-4.4 Mbp. Y-axis: each row represents a rice accession. Red: *indica* origin, including the Aus cluster. Blue: *japonica* origin, including temperate, sub-tropical and tropical clusters. Gray: unclear origin. Local ancestry analysis was performed using a set of SNP markers distributed along the whole rice genome and identified previously in 97 diverse rice accessions (Duitama et al. 2015), using custom scripts. Briefly, the accessions were first classified as *indica* or *japonica* by principal component analysis on the SNP set. Then, the most frequent SNP haplotype (MFH) was searched in the *indica* and *japonica* groups using adjacent windows of 224,000 bp (representing ∼1 cM in rice). Finally, a simple matching similarity index, SI, was calculated between each haplotype of each accession and the *indica* and *japonica* MFHs. The *indica* or *japonica* origin of the haplotype was attributed if SI > 0.99.

